# A tool for analyzing electrode tracks from slice histology

**DOI:** 10.1101/447995

**Authors:** Philip Shamash, Matteo Carandini, Kenneth Harris, Nick Steinmetz

**Author notes:** Corresponding author: Philip Shamash.

## Abstract

It is now possible to record from hundreds of neurons across multiple brain regions in a single electrophysiology experiment. An essential step in the ensuing data analysis is to assign recorded neurons to the correct brain regions. Brain regions are typically identified after the recordings by comparing images of brain slices to a reference atlas by eye. This introduces error, in particular when slices are not cut at a perfectly coronal angle or when electrode tracks span multiple slices. Here we introduce SHARP-Track, a tool to localize regions of interest and plot the brain regions they pass through. SHARP-Track offers a MATLAB user interface to explore the Allen Mouse Brain Atlas, register asymmetric slice images to the atlas using manual input, and interactively analyze electrode tracks. We find that it reduces error compared to localizing electrodes in a reference atlas by eye. See github.com/cortex-lab/allenCCF for the software and wiki.

## Introduction

Recently developed silicon probes such as the Neuropixels probe permit simultaneous recordings of hundreds to thousands of neurons across multiple brain regions (Jun et al., 2017). The resulting datasets present unprecedented opportunities to dissect neural circuit function as well as daunting challenges for data analysis (Steinmetz et al., 2018). One crucial step in the analysis is linking the recorded neurons with the brain regions where they reside. This link allows for findings to be interpreted in light of previous knowledge about that area’s function, connectivity, and cellular properties.

For probes that span multiple brain regions, however, the traditional method for identifying recorded brain regions is problematic. This method consists of labelling the electrode trajectory with a fluorescent marker or electrical lesion, slicing the brain into thin coronal sections, and comparing images of these slices to sections found in a reference atlas. This method presents three difficulties (Steinmetz et al., 2018, Luo et al., 2018). First, histology sections sliced at imperfect angles are compared to fixed-angle coronal reference images. Second, localizing regions of interest (ROIs) in the reference atlas by eye, i.e. without an image registration step, is prone to error. Third, it is not possible to visualize the trajectory of an electrode that spans multiple coronal slices.

Several tools have made progress in addressing these issues. Fürth et al., 2018 addresses the latter two problems with automated nonlinear slice registration and visualization tools but expects unangled coronal slices and a manual estimation of the anterior-posterior location. Song et al., 2018 and Xiong et al., 2018 demonstrate automated registration of angled slices but do not yet provide an interface for registration and manual fine-tuning. Methods that produce volumetric images of the brain, such as serial-section two-photon imaging and light-sheet imaging of cleared brains, can resolve all three problems (Mayerich et al., 2008, Ragan et al., 2012, Renier et al., 2014, Niedworok et al., 2016, Vandenberghe et al., 2016) but can be impractical due to the extensive setup and expensive equipment required.

Here, we address these problems with a tool for Slice Histology Alignment, Registration, and Probe-Track analysis (SHARP-Track). SHARP-Track specializes in registering variable quality brain slices cut at any angle, offering a simple MATLAB graphical user interface, and allowing users to mark distinct fluorescent tracks that may appear on the same color channel and across several slices. SHARP-Track requires brain-slice images with labelled regions of interest (e.g. fluorescent tracks) and the freely available **Allen Mouse Brain Atlas**. It uses manual user input to geometrically transform each slice image and overlay it with the reference atlas. One can then extract the coordinates of electrode tracks and plot the brain regions that they traverse.

## Methods & Results

We first describe SHARP-Track‘s registration and visualization pipeline, and then we show that it reduces error in localizing electrodes trajectories compared to marking tracks by eye.

## Registration pipeline

We developed a pipeline with a graphical user interface in MATLAB to register brain slices to the Allen Mouse Brain Atlas and identify regions of interest (Figure 1). Fist, users pre-process their histology images. This consists principally of down-sampling the images to the resolution of the reference atlas (here, 10 µm/pixel) and adjusting contrast using the MATLAB function *imcontrast* (Figure 1a). Next, by rotating a slice taken from the reference atlas, users can find the slice in the atlas matching a histological slice even if it is not perfectly coronal (Figure 1b). In the example shown, there is an angle along the medial-lateral axis of 1.8° and an angle along the dorsal-ventral axis of 0.9°. Users then click on matching points on both the reference slice and the histological slice (Figure 1b). Finally, the reference atlas and the slice image can be overlaid by geometrically transforming the slice image (Figure 1c). To accomplish this, a projective transform is generated using the MATLAB function *fitgeotrans* and the user-selected set of corresponding points between the reference atlas and the slice image. The projective transform is a linear transformation with two more degrees of freedom than the affine transform: in addition to translation, rotation, scaling, and shearing, it allows for a ‘keystone’ distortion in which parallel lines bend toward or away from each other. After refining the transformation and reference-atlas position, the map of brain regions across seemingly homogeneous brain tissue can be explored (Figure 1c). For example, the brain region where the user’s mouse is hovering can be highlighted and labelled (as in Figures 1d and 2a).

**Figure 1.**
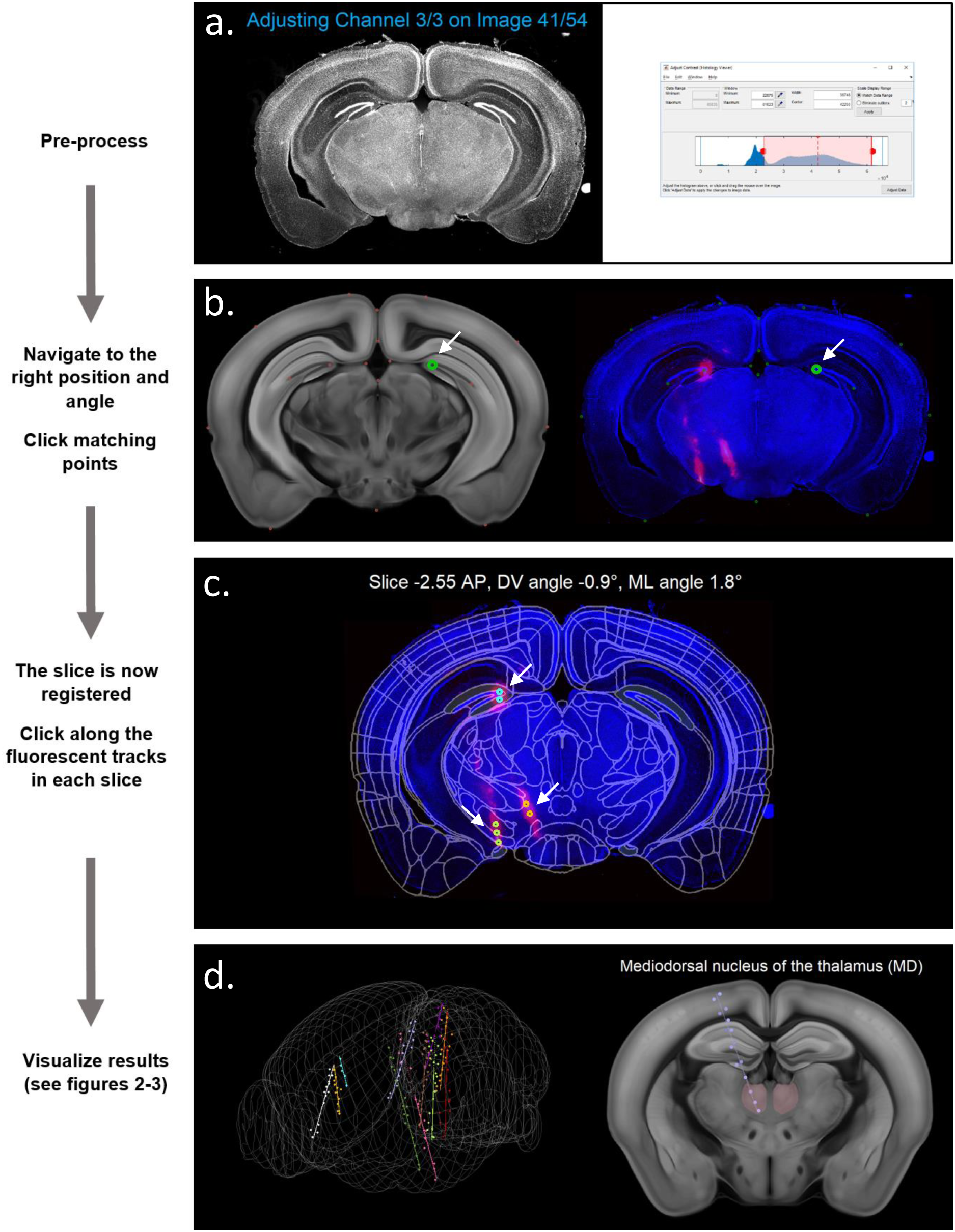
SHARP-Track’s pipeline for the manual registration of slice images and electrode tracks to the Allen Mouse Brain Atlas. **(a)** *Pre-process*. Slice images are down-sampled and pre-processed. **(b)** *Navigate.* In the GUI, users can scroll to the corresponding slice in the reference atlas. Clicking points that match between the slice image and the reference atlas then allows for image registration. White arrows point to one such pair of clicked points. **(c)** *The slice is now registered.* Brain regions present in the slice image can now be explored, and regions of interest can be marked. White arrows points to the three marked fluorescent tracks in the slice. **(d)** *Visualize results.* The trajectory of fluorescent tracks across multiple slices can be analyzed (see Figures 2-3).

Once each relevant slice has been registered in this manner, it is straightforward to scroll through these slices and click on fluorescent regions corresponding to the electrode track(s) (Figure 1c). These ROIs can then be analyzed and visualized (Figures 1d, 2 and 3). See github.com/cortex-lab/allenCCF/wiki for detailed instructions.

## Visualization of electrode trajectories that span multiple brain slices

The pipeline contains functionality for interactive visualization and verification of results (Figure 2). Here, 13 points along fluorescent tracks in 5 registered slices had been clicked, corresponding to the full track of a single electrode. The best-fit line for these points and the brain slice along which this line lies are then immediately available (Figure 2a). Best-fit lines are taken to run along the first principal component of the points and intersect their mean value. Clicking along this best-fit line brings up the point in the original histology images closest to the clicked point (Figure 2b). The estimated position, angle, and depth of probe insertion are also available (Figure 2c). Positions are reported relative to bregma; bregma was selected based on visual comparison to the Paxinos reference atlas (Franklin and Paxinos, 2012).

**Figure 2.**
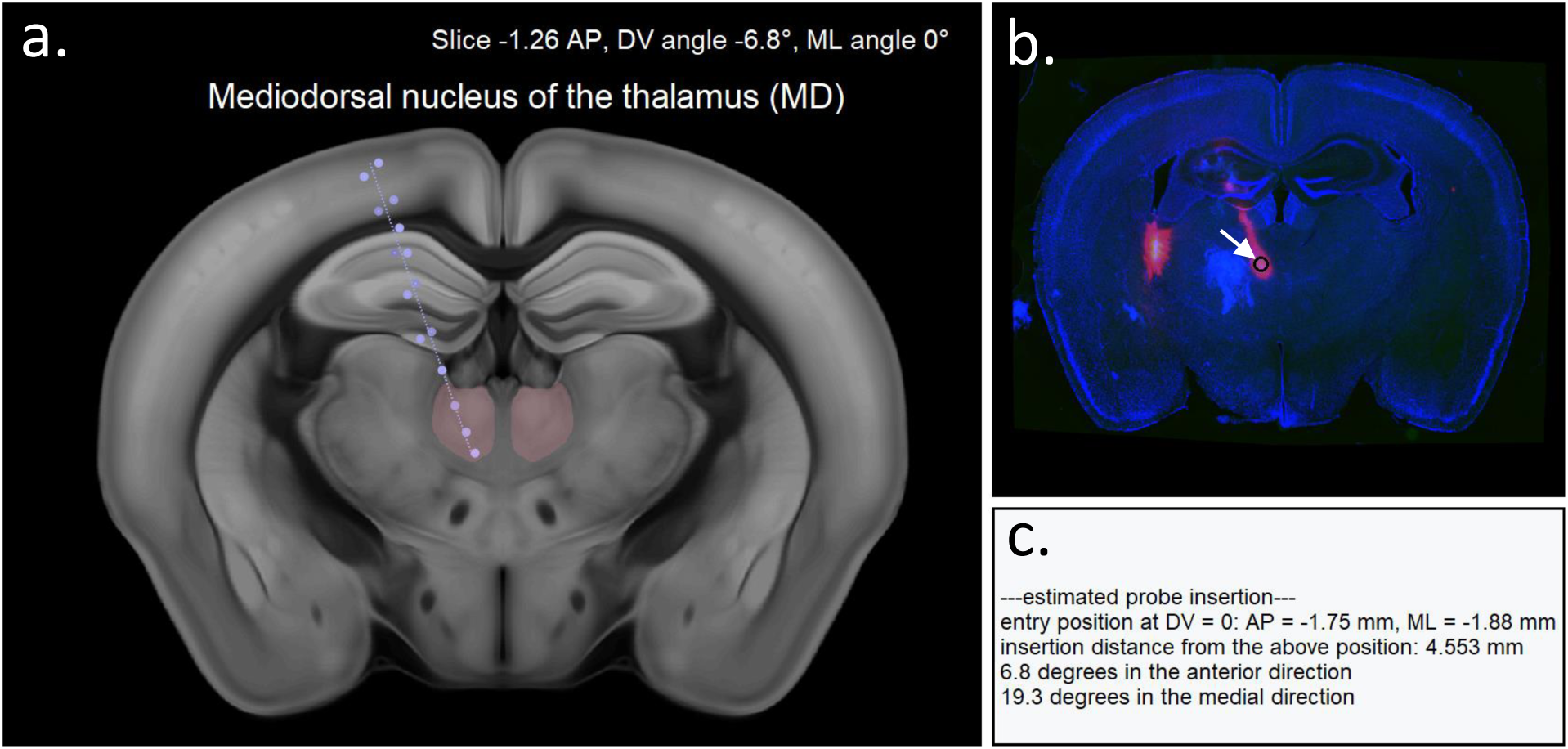
Interactive electrode-track visualization. **(a)** Circular markers are points that were clicked to mark the probe track. The best fit line for these points is displayed on the slice along which it lies (here, at a 6.8° angle on the dorsal-ventral axis), even if it spans several of the histological slices. **(b)** Clicking this track brings up the point in the histological slice images closest to the clicked point. Here, clicking the ventral MD thalamus (highlighted in panel a) brings up the image in panel b. **(c)** The estimated coordinates of the probe insertion also appear.

After the user registers slices and identifies probe track points, the software can perform an analysis of the brain regions that each track traverses (Figure 3). Here, 11 Neuropixels probe tracks marked across 44 registered slices are plotted (Figure 3a-b). In addition to a 3D rendering of the probes in the mouse brain (Figure 3a), a visualization of the brain regions lying along each probe is plotted (Figure 3bc). Brain-region labels are plotted alongside the distance from the estimated probe track to the nearest atlas annotation that differs; greater thickness indicates that the track was far from other atlas regions and can therefore be more confidently interpreted as falling within the labelled area. This value is found by sampling points in the plane perpendicular to the electrode until a distinct brain region is found.

**Figure 3.**
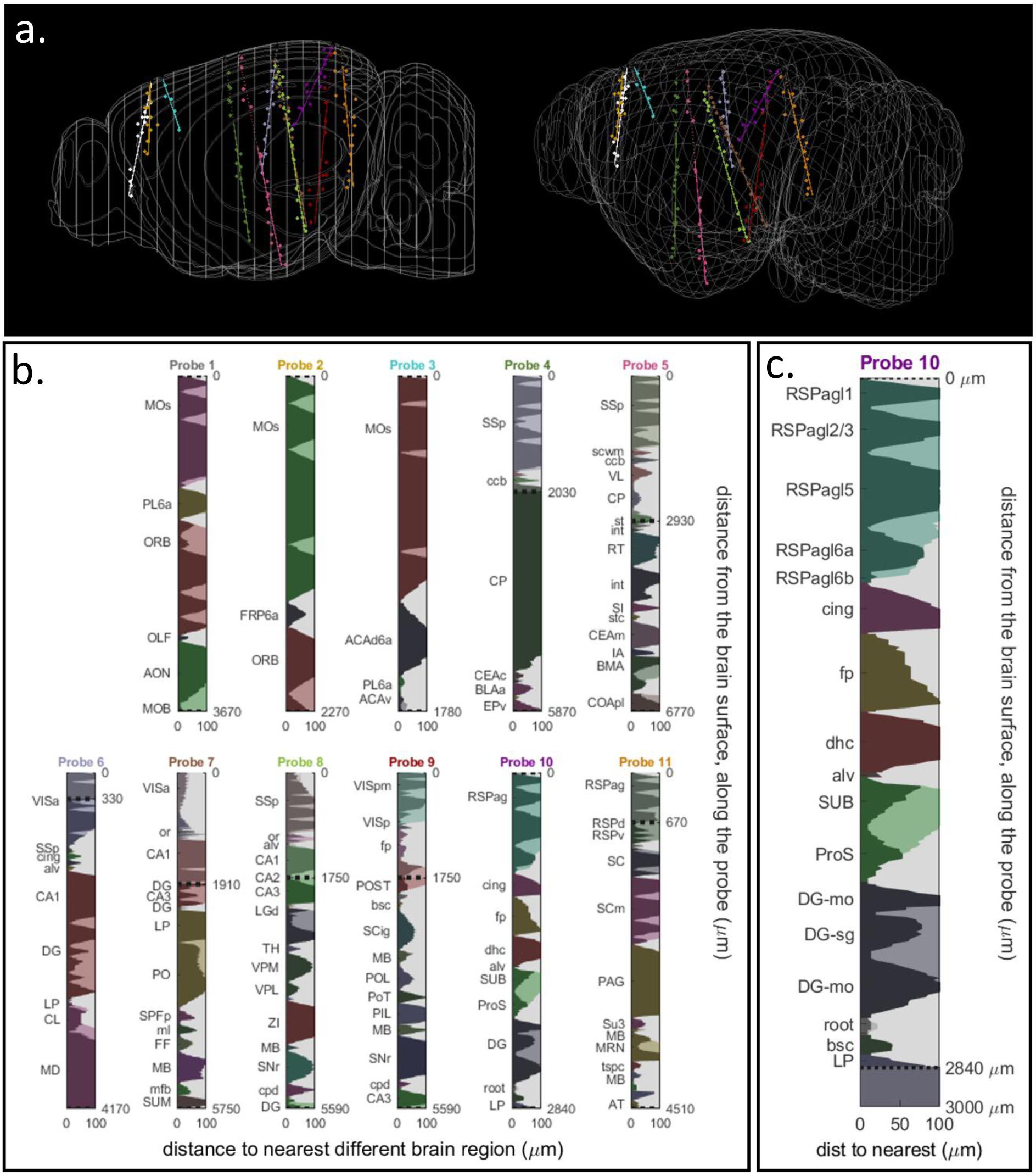
Analysis of the brain regions traversed by a probe. **(a)** The 11 probes used in this experiment were marked after registering each brain slice to the reference atlas. Best-fit lines for each electrode were then plotted in a 3D rendering of the mouse brain using SHARP-Track. **(b-c)** The brain regions lying along each probe track are also plotted. Thickness along the x-axis indicates the degree of confidence that the brain-region label is correct (see text for details). Y-axis labels indicate the identity of the brain region. Color (arbitrary hue) and shading in-between horizontal peaks indicates that regions belong to the same parent region (e.g. in panel c, the dentate gyrus molecular layer (*DG-mo*) and granule cell layer (*DG-sg*)). Dotted lines delineate the extent of the electrode’s recording channels (e.g. in panel c, at 0 and 2.84 mm). Region labels are acronyms (e.g. *RSPagl = Retrosplenial area lateral agranular part*); all acronym meanings are located in the GitHub repository in a file called *structure_tree*. Region labels are abridged in the miniature plots in panel b. **(c)** The 2.84-mm track of one electrode (the posterior-located electrode colored in purple in panel a) is plotted in full detail.

## Software and analysis

All functionalities and the graphical user interface in this paper were created with custom code in MATLAB, available from github.com/cortex-lab/allenCcf. The 3D reference atlas images and annotations come from the Allen Institute’s 10µmvoxel 2017 release using the Allen Mouse Brain Common Coordinate Framework v3, available from download.alleninstitute.org/informatics-archive/current-release/mouse_ccf or data.cortexlab.net/allenCCF

## Additional functionalities

SHARP-Track can be used to analyze non-electrode ROIs such as labelled neurons. Additionally, it can determine the entry coordinates and angle needed to target particular brain regions with a probe. See github.com/cortex-lab/allenCCF/wiki for details.

## Brain slicing and fluorescent marking

All procedures were performed at University College London and were conducted according to the UK Animals Scientific Procedures Act (1986) and under personal and project licenses released by the Home Office following appropriate ethics review. Probes were coated with DiI (Vybrant DiI Cat.No.V22885 or V22888, ThermoFisher Scientific) by holding the probe securely fixed to a rod, then drawing 2µL of DiI solution into a micropipette and slowly touching the drop of DiI to the shank of the probe repeatedly until all DiI had evaporated and dried on the shank. After this procedure, the shank was visibly pink. Probes were inserted into the brain for approximately 60 minutes, and mice were perfused and sectioned within 7 days following the first recording. Mice were perfused with 4% PFA and 60 or 90µm-thick slices were cut at approximately coronal angles. Slices were stained with DAPI and imaged with a Zeiss Axioscan at 5x magnification.

## Registering slices to the reference atlas reduces ROI localization error

We find that applying the registration step before marking electrode tracks reduces localization error. We compared the error in finding ROI coordinates from two alternative methods: 1) scrolling to the appropriate coronal slice in the reference atlas and then clicking points that appeared by eye to correspond to fluorescent regions in the histological slice image or 2) registering each slice to the reference atlas and then clicking fluorescent regions on the transformed image itself (as in Figure 1c).

First, we measured the error in finding the coordinates of ‘ground-truth’ ROIs (Figure 4a). As our ground truth, we used a **Nissl-stained brain whose entire volume had been registered to the Allen Mouse Brain Atlas** by the Allen Institute. We selected 8 2D slices from this 3D image at randomly generated anterior-posterior coordinates (with uniform probability for coordinates from +4.9 mm to −4.6 mm relative to bregma) and slicing angles (with uniform probability for angles along the dorsal-ventral and medial-lateral axes from −5° to 5°), and then overlaid 10 ROIs (red dots of 24-pixel diameter) at random positions on each slice. The same user applied the two methods to localize these ROIs. We then measured each method’s error: the distance between an ROI’s known coordinates in the Nissl volume and the coordinates extracted by the method. Registration reduced error by 42%: 218 ± 15 µm without registration vs. 126 ± 15 µm with registration, *mean*±*s.e.m*. To put these values in perspective, in a given brain hemisphere the mean volume of a brain region with its own label in the reference atlas corresponds to a sphere with a 447-µm radius.

**Figure 4.**
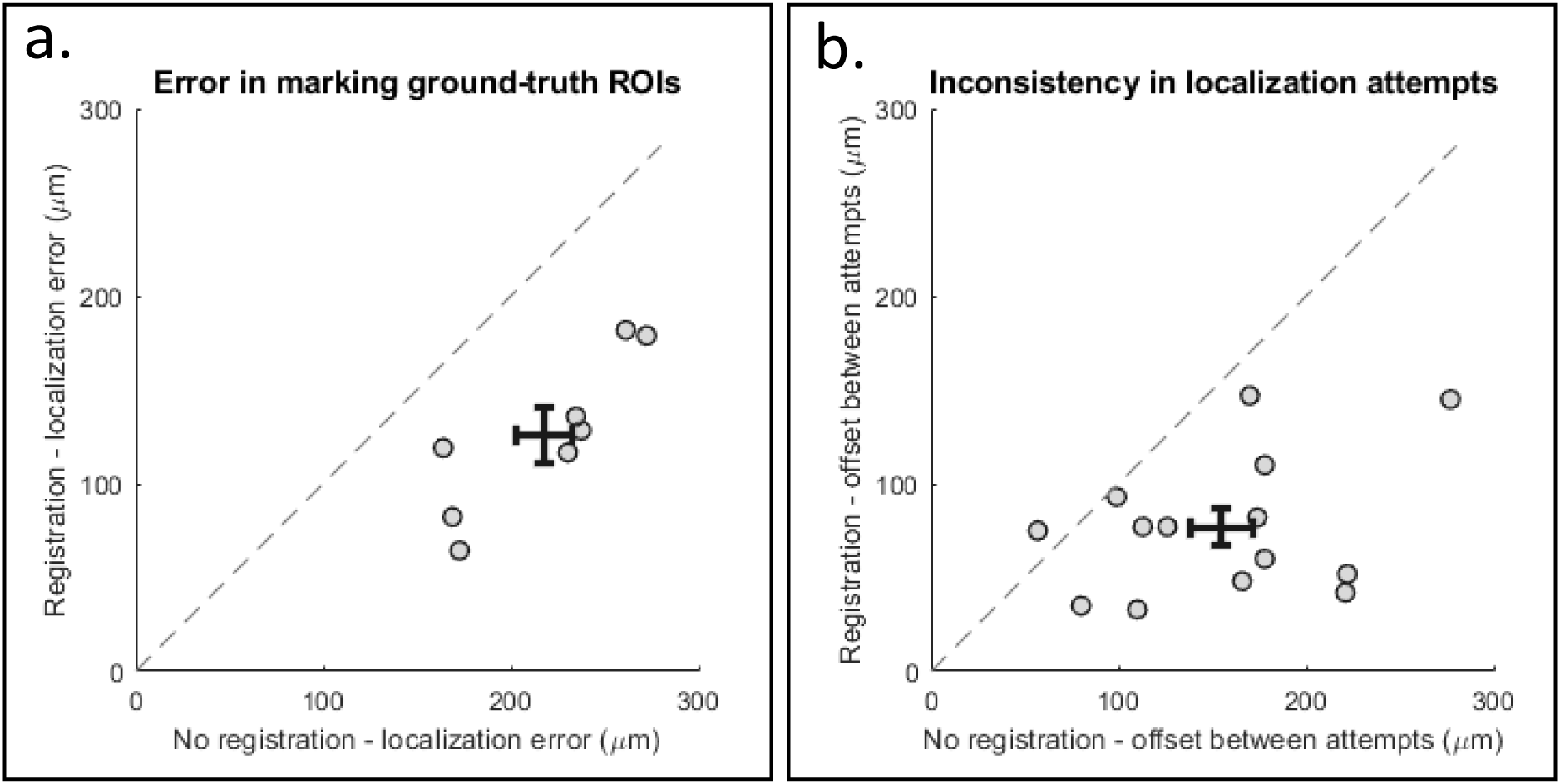
Using the registration step decreases error. **(a)** 8 slices were randomly chosen from the Allen Institute’s 3D Nissl-stained mouse brain image, and ROIs (red dots) were generated at random locations on each slice image. The two methods described in the text were used to identify the coordinates of each ROI, and then the error (i.e. Euclidean distance from the ‘ground-truth’ coordinates in the Nissl volume) was calculated. Each dot represents the average error from all ROIs in one slice. **(b)** 14 Neuropixels probe tracks from two brains were marked twice using each of the two methods described above (each probe was marked four times total). This plot shows the distance between the best-fit lines of two attempts using the same method to mark the same probe twice. It is a measure of the method’s inconsistency, and so a proxy measure of error. Each dot represents the results from one probe.

We also measured error in localizing actual electrode tracks marked with a fluorescent dye (Figure 4b). A method’s error is a combination of its bias and its variance (German et al., 1992). Since we did not have ‘ground truth’ values for these electrode trajectories, we could not measure bias. However, we did measure the variance of two probe-localization methods. Our measure of variance was: after making two attempts using a given method on a given electrode, calculating the mean distance between the two resulting best-fit lines. Distance between the lines was calculated at each dorsal-ventral coordinate from the most dorsal clicked point to the most ventral clicked point, and then averaged. This inconsistency metric is halved by using the registration step: 155 ± 16 µm without registration vs. 77 ± 10 µm with registration, *mean* ± *s.e.m.*

## Discussion

SHARP-Track offers a MATLAB user interface to explore the Allen Mouse Brain Atlas, register asymmetric slice images, and interactively analyze electrode tracks that span several slices. We find that its registration pipeline improves accuracy in localizing probes compared to marking tracks by eye.

This method has several limitations. First, creating the transforms by manual input can be time-consuming when an experiment involves many brain slices. In a future version, SHARP-Track’s manual features could be combined with an algorithm to automatically register slices such as that found in Song et al., 2018. Second, we found that there remains some error and inconsistency in marking electrode trajectories (Figure 4). We hypothesize that there are three principal reasons for this: 1) When the fluorescent dye spreads through the tissue, the best way to mark the fluorescent probe track is not obvious. 2) When selecting the position and angle that correspond to the histological slice, multiple solutions may seem correct at first. To help with this, we suggest adding a constraint: the slicing angles of consecutive slices must be similar. Additionally, applying a Nissl stain to the brain slices can make different brain slices more visually distinct. 3) Non-isometric tissue warping occurs during fixation and slicing, in particular around ventricles and edges. When this occurs, the geometrically transformed slice image will not perfectly match the reference atlas. We recommend prioritizing that the transform is accurate nearby the region of interest. An alternative approach could incorporate non-rigid transformations as in Fürth et al., 2018, Song et al., 2018, and Xiong et al., 2018.

One feature generally lacking in electrode-localization toolkits is the use of brain regions’ distinct electrophysiological signatures. A future version could allow users to tag recording channels located in an electrophysiological landmark such as white matter or ventricles. An ideal solution, however, would be the creation of an ‘electrophysiological brain atlas’ that could automatically match a channel’s activity signatures to the brain region where it was located (Jia et al., 2018,Steinmetz et al., 2018).

Finally, SHARP-Track’s interface for navigating in the Allen Mouse Brain Atlas can be applied for purposes other than post-hoc electrode localization. Initial users have found SHARP-Track useful in analyzing distributions of fluorescent cells, localizing neurons recorded with a patch pipette, and determining the parameters needed to target particular brain regions with an electrode.

## Acknowledgements

This work was funded by Sainsbury Wellcome Centre PhD Programme (PS); by postdoctoral fellowships to NS from the Human Frontier Sciences Program (LT001071/2015-L) and the European Union’s Horizon 2020 Marie Curie program (656528); by the Wellcome Trust grant to MC and KDH (grant 204915); and by the Wellcome Trust (102264, 204915), the Gatsby Charitable Foundation (GAT3531), the European Research Council (694401), and the Simons Foundation (325512). MC holds the GlaxoSmithKline / Fight for Sight Chair in Visual Neuroscience. We thank Sylvia Schröder, Daniel Regester, Dario Campagner, Rob Campbell, and Oriol Pavon Arocas for their feedback on the software and manuscript. We thank Charu Reddy, Miles Wells, Laura Funnell, and Rakesh Raghupathy for help with mouse husbandry and histology.

